# AI segmentation requires accounting for brain size to maintain performance on developmental MRI cohorts

**DOI:** 10.64898/2026.07.22.739983

**Authors:** Lena Dorfschmidt, Milly Hang Chi Mak, Sophie Adler, Konrad Wagstyl

## Abstract

The human brain undergoes rapid developmental changes through early life, underpinning the emergence of function but also marking a period of vulnerability to a range of neurodevelopmental disorders. With dynamic changes to brain size, morphology, and imaging contrast, consistent and accurate computational neuroanatomy remains a challenge. Deep learning tools for segmentation, like SynthSeg, offer robustness to heterogeneously acquired MRI contrast but remain unproven in early development. Here, we aggregated a large cohort (26k) of MRI scans spanning infant to adult development, and evaluated SynthSeg performance. Automated quality control scores, visual inspection, and spatial overlap with expert-segmented MRI scans revealed poor quality output segmentations during development. In the infant period only 36% of scans (1094/3069) passed automated QC. Rescaling infant scans to adult brain sizes significantly improved spatial overlap, and cropping scans to match adult fields of view retrieved automated quality control. Evaluation of the *SynthSeg rescale + crop* pipeline demonstrated visible and quantitative improvements in segmentation throughout infancy and childhood. There were marked increases in successful segmentations in infant scans, with 91% of scans now passing QC (2803/3069). These findings facilitate computational analysis of typical and disrupted neurodevelopment and should be considered when training the next generation of computational tools.

## 1. Introduction

During human development, the brain undergoes rapid changes in size (Bethlehem et al., 2022), shape (Bozek et al., 2018; de Vareilles et al., 2023; Mihailov et al., 2025), and tissue composition (Dubois et al., 2021), all of which complicate computational analysis of magnetic resonance imaging (MRI) in young individuals. In order to characterise neurodevelopment with adequate sample sizes, the neuroimaging community has moved towards aggregating open source datasets across overlapping age ranges (Bethlehem et al., 2022; Dorfschmidt et al., 2025; Kim et al., 2026; Sun et al., 2024). Doing so relies on the availability of imaging analysis pipelines, including for segmentation, that generalise across the human lifespan. However, developing a single pipeline to process these large datasets is difficult due to both biological and technical issues. For segmentation, contrast-agnostic deep learning tools, most notably SynthSeg (Billot et al., 2021, 2023) theoretically address the issue of changing imaging contrasts due to altered tissue composition, but because they are trained and evaluated on adult brains with adult fields of view, their performance on younger cohorts is uncertain. This study aims to quantify performance of SynthSeg across cohorts from infancy to adulthood, to identify potential reasons for failures, and to develop reliable solutions to address them.

Early post-natal development provides a particular challenge, as there are rapid changes in (i) brain size and (ii) tissue structure reflected in changes in MRI contrast. From 40 weeks gestation (i.e. full-term birth) to early adulthood, the volume of the brain increases by a factor of four to five, with different brain structures exhibiting individual trajectories. From an MRI intensity perspective, there is an inversion of the grey-white matter contrast over the first year of life as the white matter myelinates (Dubois et al., 2014). At birth, cortical grey matter appears brighter than the underlying white matter. This contrast is then lost between 4 and 12 months postnatally, before the early adult pattern emerges (Dubois et al., 2021).

In addition to these biological changes in the brain, infants are smaller than adults. For infant MRI scans, with adult-calibrated fields of view, the brain and head take up less of the image, and additional neck and shoulder anatomy is captured. This may additionally affect the generalizability of processing pipelines.

Taken together, these challenges make the segmentation of MRI scans from infants under the age of two years particularly difficult. As a consequence, multiple infant-specific processing pipelines have been developed (Hendrickson et al., 2025; Makropoulos et al., 2018; Shang et al., 2022; L. Wang et al., 2023; Zöllei et al., 2020). While age-specific pipelines may improve the individual segmentations for these specific populations, their integration into analyses over wider age windows poses additional challenges. The specific regions being segmented, how they are named, and how they are defined, often differ between pipelines. To some extent, post-hoc statistical correction can adjust for these effects, but if there are complete discontinuities (i.e. infant FreeSurfer for age <3; and adult FreeSurfer for age >3), disentangling developmental effects from methodological artifacts becomes virtually impossible.

Recent advances in artificial intelligence (AI) driven neuroimaging tools have improved our ability to automatically segment MRI scans with contrast agnostic tools (Billot et al., 2023; Gopinath et al., 2025). SynthSeg is an automated segmentation tool trained on entirely synthetic data (Billot et al., 2021, 2023). From an input segmentation, synthetic MRI data are generated with randomised contrasts, with additional data augmentations to mimic scans of differently sized brains and resolutions. Alongside robust performance across heterogenously acquired scans, SynthSeg also produces an automated, calibrated, Quality Control (QC) score. Automated flagging of poor-quality data for review or exclusion is essential for processing large cohorts where time-intensive, manual or semi-automated QC becomes unfeasible. Therefore, SynthSeg theoretically provides the opportunity to use a single pipeline for studies using aggregated datasets spanning wide age ranges, including early life.

However, the SynthSeg model was originally trained on adult-only cohorts from Alzheimer’s Disease Neuroimaging Initiative (ADNI), Open Access Series of Imaging Studies (OASIS) and the Human Connectome project (HCP), and tested on subjects with similar age distributions. Deep learning methods and particularly convolutional neural networks are known to perform poorly on samples with characteristics that are outside of their training distribution, including relatively small rescaling of objects (Azulay & Weiss, 2018). While the unique contrast features of infant scans could theoretically be captured within SynthSeg’s synthetic data generation and augmentation processes, changes in brain size are not. In particular, the total cerebral volume of newborn infants is approximately 25% of a typical adult used to train the SynthSeg model. These differences in morphology and size of infant brains highlight a significant need to evaluate SynthSeg across a broad range of neurodevelopment including early infant scans.

Here, we set out to assess SynthSeg’s performance from birth to adulthood, and to develop an adjustment to the tool that would allow researchers to use a single pipeline across this age range. To this end, we made use of two cohorts: (i) We curated a large aggregated MRI dataset of 16 open source studies, jointly spanning the age range from 20 weeks post-menstrual age to 35 years of age - the “Neurodevelopmental cohort”, allowing us to assess SynthSeg’s performance across a wide range of ages and cohorts; (ii) We used the Baby Open Brains (BOBs) Repository MRI data, an open source dataset, which includes expert-curated ground truth segmentations for infants aged 1-9 months. This BOBs dataset enabled us to make direct comparisons between ground truth segmentations and SynthSeg-generated segmentations and QC scores.

Objective 1: Assess SynthSeg’s performance throughout the paediatric and early adulthood period. (i) First, we assessed changes in SynthSeg’s automated QC measures. We hypothesized that SynthSeg would perform worse in infant scans. (ii) To assess whether SynthSeg’s age-related changes in automated QC estimates translate into actual decreases in segmentation accuracy, we used the BOBs data to assess the spatial overlap of SynthSeg-generated and ground truth segmentations.

Objective 2: We hypothesized that SynthSeg’s performance may be related to brain size. Therefore, it may be possible to improve the pipeline’s performance by transforming infant MRI data to appear more similar to the data on which SynthSeg was trained. Specifically, we hypothesized that age-specific rescaling of infant MRI scans to approximate adult volumes and then cropping their field-of-view to match adult MRI would (i) improve the spatial overlap with ground truth segmentations; (ii) lead to correspondingly improved automated QC scores; and (iii) enable a significantly higher proportion of scans across neurodevelopment to be segmented and volumes extracted for downstream analysis.

## 2. Methods

### 2.1 Neurodevelopmental Datasets

In accordance with King’s College London ethics guidance, secondary analysis of anonymised datasets does not require explicit ethical approval. We aggregated 16 open source MRI datasets, jointly spanning the age range from 20 weeks post menstrual age to 35 years to form a large aggregated neurodevelopmental cohort. These data include: the Developing Human Connectome Project study (dHCP; *N_subjects_* = 600; 20-44 weeks post conception; (A. D. Edwards et al., 2022; D. Edwards et al., 2021)), the HEALthy Brain and Child Development study (HBCD; *N_subjects_* = 602; 39-78 weeks gestational age; (Chambers et al., 2024; Nelson et al., 2024)), the Baby Connectome Project data; (BCP; *N_subjects_* = 372; 0-7y; (Howell et al., 2019)), the University of California San Diego Biomarkers of Autism at 12 Months Study (UCSD; *N_subjects_* = 324; 1-7y; (of Excellence (ACE) Network, 2008)), the Infant Brain Imaging Study (IBIS; *N_subjects_* = 433; 0-3y; (Piven & of Excellence (ACE) Network, 2008)), the NIH Pediatric MRI Release (NIHPD; *N_subjects_* = 554; 0-22y; (Evans & Group, 2006; Rivkin et al., 2010)), Calgary (*N_subjects_* = 126; 1.9-8y; (Reynolds et al., 2019)), the OpenNeuro-Pixar study( *N_subjects_* = 155; 3.5-39y; (Richardson et al., 2018a, 2018b)), the OpenNeuro-Wang study ( *N_subjects_* = 322; 5-9y; (J. Wang et al., 2022a, 2022b)), the Brazilian High Risk Cohort study (BHRC; *N_subjects_* = 610; 5-14y; (Brazilian High Risk Cohort Study (BHRC), 2009; Salum et al., 2015)), the Human Connectome Project Development data (HCP-D; *N_subjects_* = 652; 5-21y; (Harms et al., 2018; Somerville et al., 2018, 2020), the Developing Chinese Connectome Project data (devCCNP; *N_subjects_* = 195; 6-18y; (S. Liu et al., 2021)), the Philadelphia Neurodevelopmental Cohort (PNC; *N_subjects_* = 1601; 8-23y; (Satterthwaite et al., 2014, 2016)), the Adolescent Brain Cognitive Development study (ABCD; *N_subjects_* = 11, 865; 9-14y; (Casey et al., 2018; Jernigan et al., 2025)), the Neuroscience in Psychiatry Network study (NSPN; *N_subjects_* = 316; 14-26y; (Kiddle et al., 2018; Whitaker et al., 2016)]), the Human Connectome Project Young Adult data (HCP; *N_subjects_* = 1206; 22-37y; (Van Essen et al., 2012)). This sample served as a means to assess pipeline performance over a broad neurodevelopmental period. All subjects and sessions required a T1w or T2w 3D anatomical MRI scan, age at MRI scan and sex information for inclusion. Age at MRI scan was calculated as post-menstrual age in days. If both T1w and T2w data were available, the T1w was used. If multiple runs were available for a session, 1 run was selected. Subjects were excluded for the following reasons: 1) Age or sex information was unavailable or incorrectly coded; 2) Subject had a diagnosis of a neurodevelopmental disorder; 3) MRI data did not have full field of view or 4) MRI data file in an unusable file format.

### 2.2 Expert labelled sample

The second dataset used in this study is a subset of the BCP data, for which manually segmented ’ground truth’ segmentations exist. The Baby Open Brains (BOBs) Repository MRI data (Feczko et al., 2024) is an open source dataset of manually curated and expert-reviewed manual infant brain segmentations. The dataset includes anatomical T1- and T2-weighted MRI data from 71 imaging sessions across 51 infant participants. Participants were scanned between the ages of 1 and 9 months. We found 62 matched sessions across BCP and BOBs. These matched sessions were used to directly compare SynthSeg-generated segmentations with manually curated “ground truth” labels.

### 2.3 SynthSeg performance across paediatric development

To assess SynthSeg performance across neurodevelopment, we processed the neurodevelopmental cohort and the expert-labelled sample using SynthSeg (Billot et al., 2021, 2023) with the ‘--robust’ flag. This resulted in SynthSeg-generated segmentations (**Fig. 1**), as well as pipeline-supplied automated quality control (QC) scores for eight gross anatomical structures: general grey matter, general white matter, the brainstem, the thalamus, pallidum+putamen, the cerebellum, and hippocampus+amygdala, as well as the CSF. The per-structure QC scores are generated by a regression network, originally trained to predict the DICE score of the output segmentation for each of these structures. Following the SynthSeg developer recommendations, the summary QC score was the minimum QC score across structures for each subject and a threshold of 0.65 was used to classify segmentations as being of adequate (pass) or inadequate (fail) quality (**Fig. 1A-C**).

**Fig. 1:**
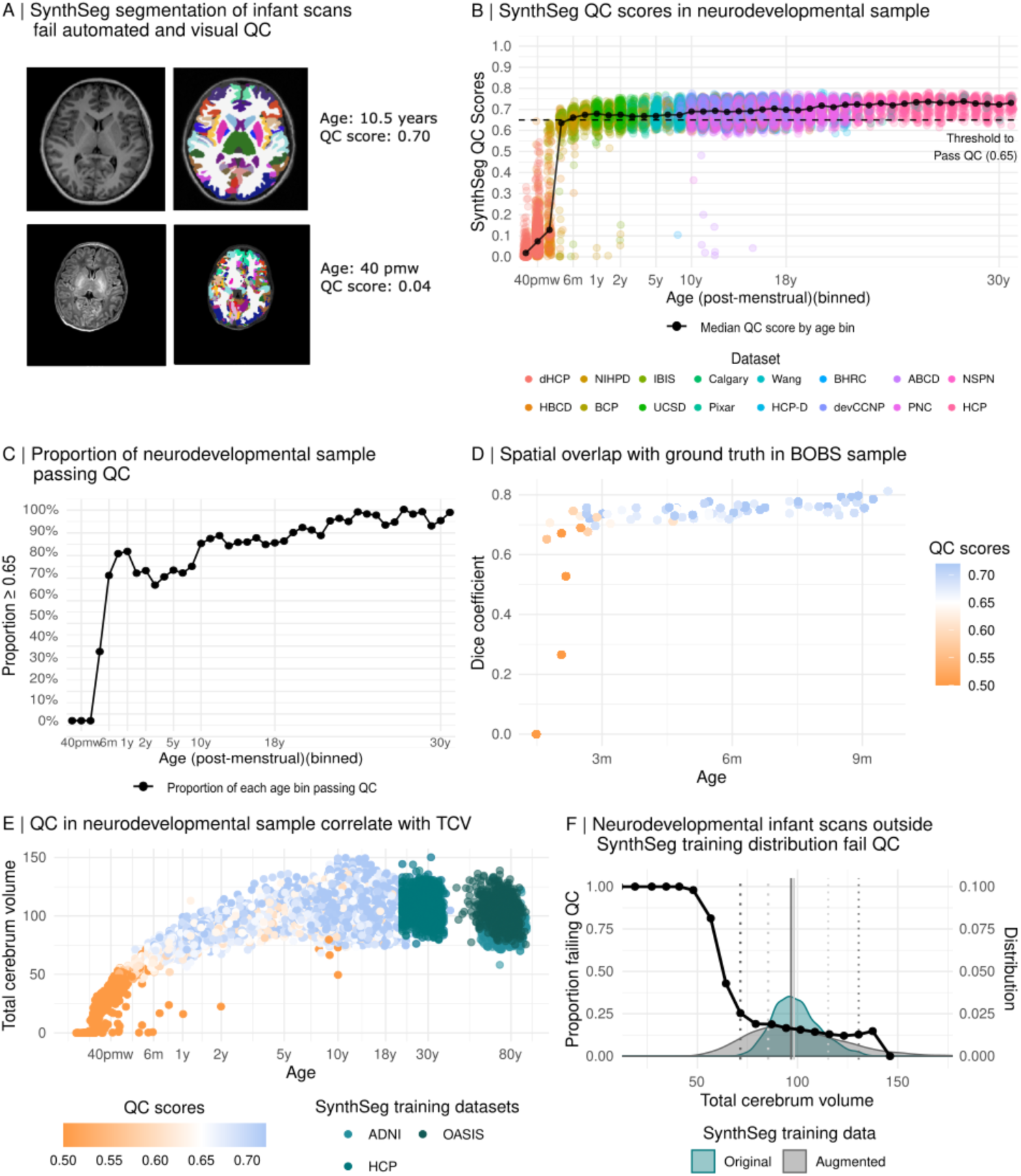
SynthSeg automated QC metrics are highly dependent on age at scan. **(A)** Left T1w MRI of 10 year old and neonate, exhibiting characteristic differences in MRI contrast and brain size. Visual inspection of SynthSeg outputs (right) reveals good quality segmentation at age 10, but poor in the infant, accurately reflected in the SynthSeg-generated automated QC score. **(B)** SynthSeg QC metrics increase with age, particularly below 2 years of age in an aggregate sample of 18,000 T1w and T2w MRI scans. Each scan is a circle, and is coloured according to the dataset of origin. The black line indicates the median QC score of each age bin across datasets. **(C)** The percentage of scans in each age bin that pass automated quality control (QC ≥ 0.65). Very few infant scans pass QC, and the percentage of passing scans increases as age increases, a trend that continues into adulthood. **(D)** In an expert-labelled infant MRI sample (BOBs), the spatial overlap between SynthSeg segmentations and ground truth labels (estimated using Dice score) is systematically lower in younger subjects, where QC scores (colour of circle) are also lower. **(E)** In the neurodevelopmental cohort, Total Cerebrum Volume / 1000 mm^3^ (TCV) is plotted against age and coloured by the minimum SynthSeg QC value. Subjects with lower TCV have systematically lower QC scores. In the three synthetic training datasets that SynthSeg was trained on (green dots), the TCV and age do not overlap our neurodevelopmental sample. **(F)** Distributions of TCV of the Synthseg training data are plotted with and without random augmentation. Solid vertical lines indicate the median of the original (light grey) and augmented (dark grey) distributions, and the dotted lines indicate the 5th and 95th quantiles. The proportion of scans in the neurodevelopmental sample that failed QC is overlaid (black line). As TCV decreases and falls outside the Synthseg training data TCV distribution the proportion of scans failing QC increases.

### 2.4 Evaluation of SynthSeg training data

SynthSeg is trained on synthetic data generated from a subset of subjects from three studies: the Human Connectome Project Young Adult (HCP; (Van Essen et al., 2012)) data, the Alzheimer’s Disease Neuroimaging Initiative study (ADNI; (Jack et al., 2008)) and the Open Access Series of Imaging Studies data (OASIS; (Marcus et al., 2007)). However the subject IDs of the specific subset used to train SynthSeg are not available. To estimate the approximate distribution of brain sizes within data used for SynthSeg training, we extracted the measurements for total cerebral volume for these 3 cohorts. Total cranial volume estimates were calculated using SynthSeg robust for ADNI and HCP, and preprocessed Freesurfer total cranial volume for OASIS.

To develop the original SynthSeg model, these training data were augmented using random spatial transformations, including random rescaling (up to +/-20%) of the images. To approximate the impact of this rescaling on the distribution of training volumes, in line with SynthSeg, for each individual scan, we randomly sampled a linear scaling factor for each image dimension [0.8,1.2] and rescaled the total intracerebral volume accordingly, generating 10,000 such samples. The unaugmented (original) and rescaled (augmented) total cranial volume estimates for the SynthSeg training cohort were compared with QC metrics and SynthSeg volume estimates for the Neurodevelopmental cohort (**Fig. 1E**&**F**).

### 2.5 MRI preprocessing pipelines

We hypothesized that SynthSeg performs better on (1) brains with sizes represented within their training distribution and (2) cropped to have similar fields of view. We therefore tested the performance of four pipelines: (i) Unaltered SynthSeg: “*SynthSeg original”*; (ii) SynthSeg on cropped MRI scans: “*SynthSeg cropped”*; (iii) SynthSeg on MRI scans that have been rescaled to approximately adult size: “*SynthSeg rescaled”*; (iv) SynthSeg on MRI scans that have been rescaled to approximately adult size and subsequently cropped: “*SynthSeg rescale+crop”* (**Fig. 2**). The respective rescaling and cropping steps are described in detail below:

**Fig. 2:**
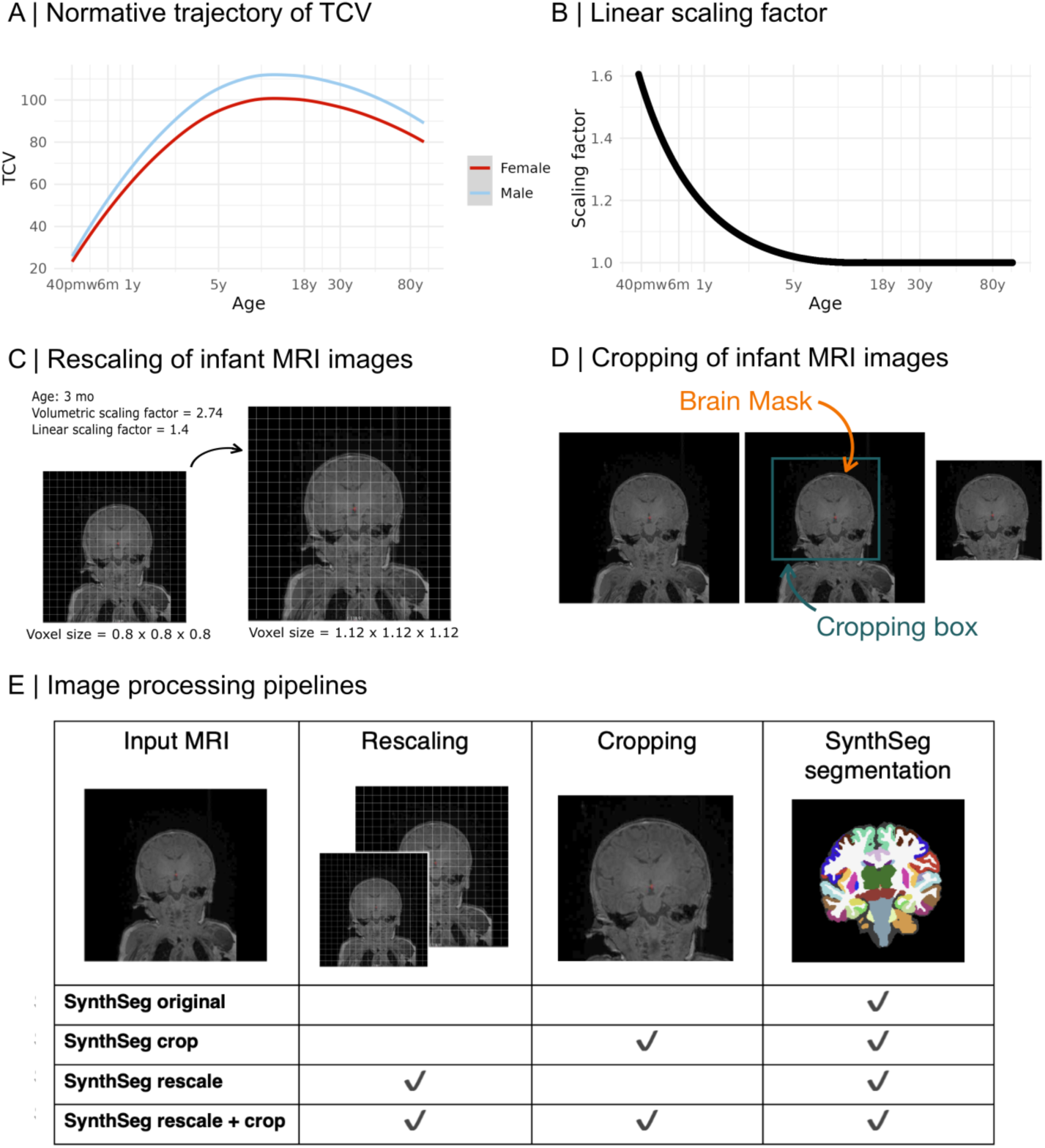
Pipeline development. **(A)** Prior work has shown large changes in total cerebral volume (TCV) across the lifespan (Bethlehem et al. 2022). For example, at six months of age, the median normative infant TCV is about 80% that of the healthy population peak volume reached around 12 years of age. **(B)** We used prior trajectories to estimate the scaling factor of infant brains to the maximum lifespan volume. The rapid increase in TCV in early life is reflected in a pronounced drop in the scaling factor until about 8 years of age. Since SynthSeg does not usually have difficulties processing scans from the end of life, in this work, we round the scaling factor to 1 for individuals above 15 years of age. **We tested the effects of two preprocessing steps on SynthSeg performance. (C)** Rescaling: We rescaled MRI scans to be of roughly adult size. Specifically, we changed the voxel-size in the image header using the above age-based scaling factor. **(D)** Cropping: we cropped scans to reproduce the approximate adult field of view in infant scans. We ran SynthStrip to derive a brain mask, and cropped each image in a 25mm box around the brain mask. **(E)** We compared the segmentation results of four pipelines: (i) SynthSeg original; (ii) SynthSeg + cropping; (iii) SynthSeg + resizing; and (iv) SynthSeg rescale + crop.

### 2.6 Image rescaling

We rescaled infant and paediatric MRI scans to be approximately adult size (**Fig. 2A**). To this end, for each scan, we derived an age-specific scaling factor based on a previously published normative trajectory of total cerebral volume (TCV) across the lifespan (Bethlehem et al., 2022). We downloaded the models and code published in: https://github.com/brainchart/Lifespan. We then estimated the percentage of the maximum TCV that a normative infant scan at a given age would have relative to the maximal TCV observed across the lifespan (**Fig. 2C)**. This scaling factor (**Fig. 2D**) was then applied to adjust the resolution parameters in the MRI scan header prior to running the image through SynthSeg. SynthSeg rescaling augmentations on adult capture within-adult brain size variability. Thus, the scaling factor is limited to 1 (i.e. no rescaling) for individuals whose age is at or above the age of peak TCV.

### 2.7 Image cropping

For a given MRI scan, we used SynthStrip(Hoopes et al., 2022) to derive an approximate brain mask (**Fig. 2B**). By extending 25mm beyond the brain mask, we then created a cropping box. For rescaled scans, this distance was estimated in rescaled-space. We cropped each scan in all six image directions.

### 2.8 Spatial overlap with ground truth

We used the expert-labelled sample to assess the quality of the segmentations for each of the pipelines by calculating the spatial overlap between the SynthSeg-predicted and ground truth manually segmented brains. First, we registered the BOBs manually-curated segmentations into BCP-space. More specifically, we registered subjects’ BOBs T1-weighted images to the BCP T1w and T2w images using a rigid registration in ANTsPy (Klein et al., 2009). We then applied the transform to the BOBs segmentations using a nearest-neighbor approach. We visually assessed and quality checked the segmentations overlaid on the BCP scans.

We assessed the spatial overlap between the SynthSeg-based and ground-truth segmentations in two ways: (1) SynthSeg automated quality-control is performed in broad tissue classes. To match this approach, we merged the SynthSeg segmentation labels to match the tissue classes used in the QC score estimates, i.e. we generated labels for: general grey matter, general white matter, cerebellum, brainstem, thalamus, putamen+pallidum, and hippocampus+amygdala. CSF was not segmented in BOBS and was excluded from this analysis. (2) We subsequently applied a more fine-grained approach, where we assessed the spatial overlap between regional labels. However, the BOBs segmentations include labels for cerebral gray and white matter and 23 subcortical structures, whereas SynthSeg segmentations include regional cortical labels. We therefore merged the SynthSeg segmentation labels to match the BOBS labels for regional comparisons in the subcortex, as well as the two broad cortical classes.

We then estimated the Dice-coefficient of each SynthSeg-generated label with their corresponding manually-generated counterpart as:

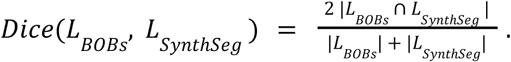

Where *L_BOBs_* is the expert-defined label from BOBs, and *L_Synt_*_ℎ*Seg*_ is the same label from the SynthSeg segmentation.

The *SynthSeg original* and *Synthseg rescale+crop* pipeline were also applied to all T2w images in the BOBs cohort.

## 3. Results

### 3.1 SynthSeg performance during paediatric development

We assessed SynthSeg’s segmentation performance during the neurodevelopment period. To this end, we processed the large aggregated sample of 26,716 MRI scans (25,911 T1w, 805 T2w) included in our neurodevelopmental cohort using SynthSeg with the ‘--robust’ flag. We found 0% (0/1493) of infant scans below 3 months, and only 32.72% (71/217) of those between 3-6 months, passed automated QC (**Fig. 1A&B**). Furthermore, the percentage of scans that passed QC continued to increase until at least 10 years of age (**Fig. 1C**). Multiple independent datasets with overlapping age groups were included in this analysis, such that particular acquisition sequences or scanners were unlikely to be the underlying cause (**Fig. 1B**). In the BOBs dataset, for which there was expert-annotated ground truth, the same relationship was present, with Dice coefficient significantly decreasing with decreasing age (ρ=0.73, p<0.001; **Fig 1D**).

We hypothesized that automated quality control metrics were strongly related to brain size. To this end, we assessed the relationship between QC scores and total cerebrum volume (TCV). We observe a clear relationship, with smaller brains showing dramatically decreased QC scores (ρ =0.32, p<0.001; **Fig. 1E**). This is particularly pronounced in the 0-2 year age range (ρ =0.88, p<0.001).

To better understand the relationship between age, brain size and QC scores, we compared the TCV of the neurodevelopmental cohort with the three adult cohorts used to train SynthSeg: HCP, ADNI and OASIS (**Fig. 1E**). We found that the distribution of raw TCV values in these three cohorts does not cover the entire distribution of TCV in the neurodevelopmental cohort (**Fig. 1E**).

During training, SynthSeg increases variation within the dataset using augmentations including random spatial transforms. Importantly, for our purposes, this includes rescaling each voxel dimension independently using a random scaling factor sampled from the uniform distribution [0.8, 1.2], aimed at improving model robustness to differences in brain size. To estimate the effects of this rescaling on the brain size distribution of the SynthSeg training data, we sampled this random rescaling 10,000 times and applied the combined factors to the TCV estimates.

These transformations widen the distribution of TCV values in the synthetic data, but the distribution still does not cover the range observed in early development (**Fig. 1F**). The QC failure rate appears to closely track this distribution, increasing dramatically as subjects TCVs begin to be less well represented in the training distribution (**Fig. 1F**).

### 3.2 Comparison of pipelines

We hypothesized that SynthSeg would produce higher quality segmentations with well-calibrated automated QC scores on infant scans that were rescaled to adult brain sizes **(Fig. 2B&C)** and cropped to a matching field of view **(Fig. 2D)**.

We first tested the following three different pipelines **(Fig. 2E)** on each infant scan of the BOBs sample, to determine the pipeline that ensured best agreement with the ground truth segmentations: (i) *SynthSeg original*, (ii) *SynthSeg crop*, (iii) *SynthSeg rescale*, (iv) *SynthSeg rescale+crop.* 57/62 subjects successfully passed through all pipelines. Five subjects were excluded either due to poor registration of BOBs segmentation to the BCP-provided image (1/5) or due to memory errors in the rescale-only pipeline (4/5).

We estimated the spatial overlap between automated and ground truth segmentations as the Dice score between model outputs and BOBs manually curated segmentations. Using the minimum Dice score across regions as a summary measure of segmentation performance, we found that resampling pipelines led to significant increases in Dice score relative to the *SynthSeg origina*l and *+crop* pipelines (*t*(56)=3.64, *p*_Bonferroni_=0.004; *t*(56)=3.54, *p*_Bonferroni_=0.005, **Table 1**; **Fig. 3A**, **C**). The largest improvements in Dice coefficient were seen in cortical grey and white-matter segmentations (**Fig. 3F; Tables S1 & S2**).

**Fig. 3:**
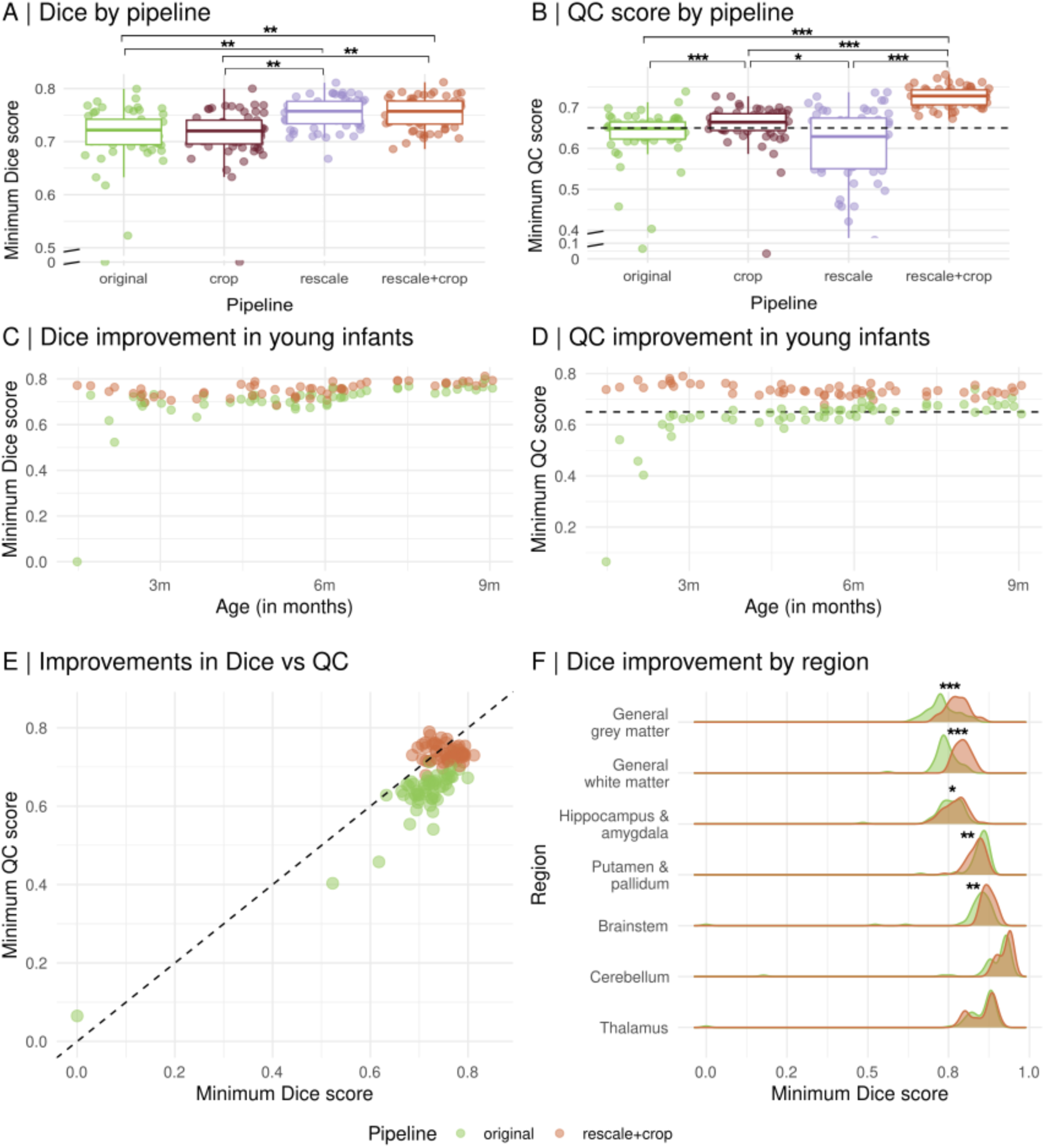
Improvements in spatial overlap and automated QC in expert-labelled segmentations. We computed the Dice score **(A)** and automated QC score **(B)** for outputs from *SynthSeg original, crop, rescale and rescale+crop,* using 57 expertly labelled scans in the BOBs dataset. Pipelines with rescaling had significantly higher Dice scores, but only *rescale+crop* also exhibited significant corresponding increases in the QC scores. Age-related changes in Dice **(C)** and QC **(D)** scores were seen in *SynthSeg original* but not *SynthSeg rescale+crop* pipelines, with all BOBs samples above the recommended minimum QC threshold of 0.65 (dashed black line). **(E)** SynthSeg’s QC score is trained to estimate minimum Dice score across broad tissue classes. Improvements in minimum tissue Dice score agreement with expert labels were reflected in corresponding increases in automated QC scores. **(F)** Region-level Dice score comparison between *SynthSeg original* and *rescale+crop* pipelines. *SynthSeg rescale + crop* improvements were largest in the general (cortical) grey and white matter, with a small decrease in “Putamen & pallidum”, driven specifically by the putamen (Fig S1).

**Table 1:** Pairwise comparisons of performance between pipelines. Dice and QC scores compared between each pipeline using paired t-tests and corrected for multiple comparisons using Bonferroni. Significant p-values after correction are indicated by stars (* p < 0.05, ** p < 0.01, *** p < 0.001). *Rescale+crop* performs significantly better than the *original* pipeline in both Dice (p=0.004) and QC score (p<0.001) measurements.

| Pipelines |  | Dice Score |  |  |  | QC Score |  |  |  |
| --- | --- | --- | --- | --- | --- | --- | --- | --- | --- |
| Pipeline 1 | Pipeline 2 | Mean dice difference | t-statistic | p-value | Bonferroni p-value | Mean QC difference | t-statistic | p-value | Bonferroni p-value |
| original | crop | 0.0027 | 0.9 | 0.373 | 1 | 0.023 | 5.58 | 7.20E-07 | 4.30E-06*** |
| original | rescale | 0.0495 | 3.64 | 6.00E-04 | 0.004** | -0.0184 | -1.37 | 0.177 | 1 |
| **original** | **rescale+crop** | **0.0495** | **3.61** | **6.60E-04** | **0.004\*\*** | **0.0975** | **7.73** | **2.20E-10** | **1.30E-09\*\*\*** |
| crop | rescale | 0.0468 | 3.57 | 7.40E-04 | 0.004** | -0.0414 | -2.96 | 0.004 | 0.027* |
| crop | rescale+crop | 0.0468 | 3.54 | 8.10E-04 | 0.005** | 0.0744 | 6.24 | 6.10E-08 | 3.60E-07*** |
| rescale | rescale+crop | 0 | -0.05 | 0.963 | 1 | 0.1158 | 9.34 | 5.00E-13 | 3.00E-12*** |

The ability to generate reliable automated quality control scores is invaluable when working on large MRI samples. Thus, we next assessed whether the improvement in the spatial overlap between ground truth and automated segmentation quality translated into corresponding improvements in SynthSeg-generated automated QC scores. Automated QC scores indicated sub-threshold (QC<0.65) segmentation quality for 50.9% of the BOBs subjects with the *SynthSeg original* pipeline. In comparison, 100% in the *SynthSeg rescale+crop* pipeline passed QC (**Fig. 3B**), reflecting the improvements in Dice overlap (**Fig. 3D&E**). Interestingly, while both *rescale* and *rescale+crop* pipelines led to improvements in Dice overlap between SynthSeg and expert segmentations, corresponding improvements in QC score were only seen in the *SynthSeg rescale+crop* pipeline (**Fig 3B**).

Similar improvements in Dice and QC scores were found in the *SynthSeg rescale+crop* pipeline relative to *SynthSeg original* when applied to the T2w BOBs MRI scans **Fig. S1.**

### 3.3 Pipeline performance over the paediatric age range

Lastly, we assessed whether the improvements in segmentation performance in the BOPs sample, using the proposed *SynthSeg rescale+crop* pipeline, extended to the wider neurodevelopmental period and across samples with different imaging protocols and acquisition qualities. We applied the *SynthSeg rescale+crop* pipeline to the full neurodevelopmental cohort, and found there were improved QC scores compared to the original SynthSeg processing, in particular in subjects under two years (**Fig. 4A,B&C**). Across all age bins 87% of scans (23205/26706) passed QC using *SynthSeg rescale+crop,* compared to 79% using *SynthSeg original.* For ages 0 to 2 years, where only 36% of scans (1094/3069) passed QC using *SynthSeg original*, 91% of scans passed QC for *SynthSeg rescale+crop* (2803/3069). Visual inspection of selected scans showed visible improvements in segmentation quality for scans now passing QC **(Fig. 4A)**, with low QC scores continuing to flag remaining poor quality segmentations (**Fig. S2**).

**Fig. 4:**
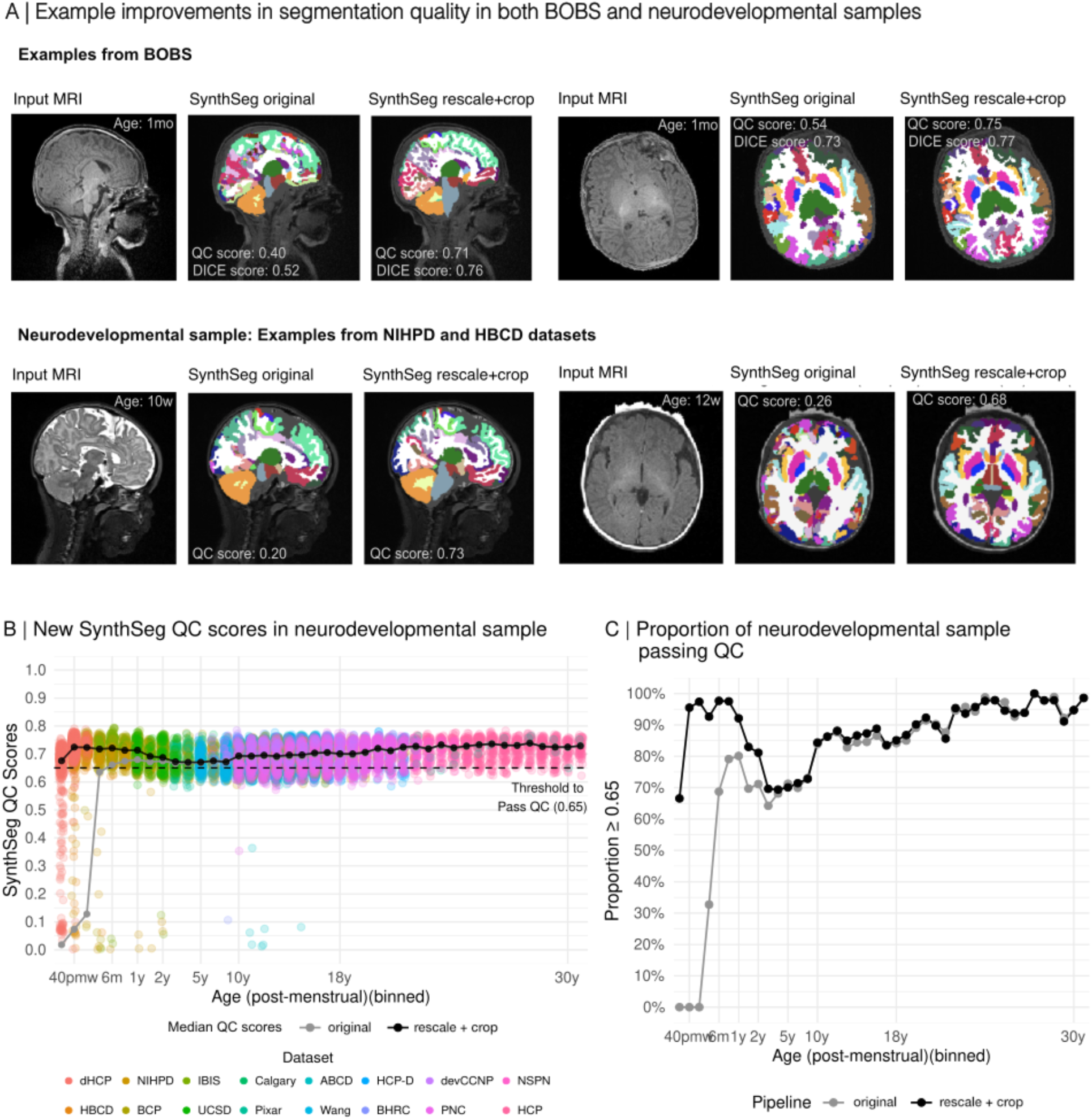
Improvement in segmentation performance across cohorts. **(A)** Four example individuals that failed automated QC using *SynthSeg original*, but passed QC using *SynthSeg rescale + crop*. Reflecting these QC scores, segmentations exhibit clear improvements in quality. The BOBs cohort subjects have corresponding expert labels, allowing calculation of corresponding Dice scores. **(B)** QC scores using *SynthSeg rescale + crop* in the neurodevelopmental cohort. The median QC scores with the new *SynthSeg rescale + crop* (black line) are above the QC threshold for all age groups, which is not the case for *SynthSeg original* (grey line). Each scan processed through the new *SynthSeg rescale + crop* pipeline is a circle, and is coloured according to the dataset of origin. **(C)** The proportion of scans which are above QC threshold of 0.65 for *SynthSeg rescale + crop (i.e. pass)* and *SynthSeg original* (black). 91% of *rescale + crop* scans aged 0-2 passed QC compared to 36% for the *original* pipeline.

## 4. Discussion

We propose an adapted *SynthSeg rescale+crop* pipeline for automated segmentation of brain MRI scans from infants to adulthood. Evaluation of the original SynthSeg pipeline demonstrated high failure rates in infant scans which were related to smaller brain sizes. Hypothesising that this was primarily due to infant brain sizes being outside the original SynthSeg adult-only training distribution, we demonstrated that rescaling and cropping scans, prior to SynthSeg inference, significantly increased both visual and quantitative segmentation quality, along with corresponding improvements in the automated QC scores. The *SynthSeg rescale+crop* pipeline is made openly available for reuse.

*SynthSeg rescale+crop* preserves two invaluable features of SynthSeg; high quality contrast-agnostic volumetric segmentation and quality control metrics that enable automated identification and removal of poor-quality segmentations. Initially, we identified that infant brain sizes fell outside of the training cohort distribution and that by rescaling brains to the volumes of adult brains, the segmentation Dice scores were significantly improved on a gold-standard segmented cohort (Feczko et al., 2024). SynthSeg’s QC network is designed to predict the quality of the segmentation (Dice score) for major tissue classes. However, while rescaling alone increased Dice scores, automated QC scores counterintuitively decreased. If relying on the automated QC scores when analysing large cohorts, using rescaling alone would lead to the incorrect discarding of scans with high-quality segmentations. The QC prediction neural network includes a final global averaging step across neurons with differing fields of view. Infant MRI scans have a relatively larger anatomical field-of-view than adults, capturing more space around the brain. Infant MRIs therefore have larger patches of the MRI scan without tissue to segment, where the network is likely predicting low Dice scores. Aggregating these areas into the final QC scores likely depresses QC scores despite high-quality segmentation where the brain tissue is present. Cropping prior to SynthSeg segmentation and QC prediction allows the QC scores to capture the increased segmentation quality and results in improvements in Dice scores being reflected in improved QC scores in the BOBs dataset, whilst still producing low QC scores on segmentations of poor quality (**Fig. 3 A,B & S2**)

SynthSeg, through synthetic data generation and augmentation processes, was developed to be contrast agnostic. Consistent with this, we demonstrate that similar improvements in performance were seen in T2w infant MRI scans, albeit with a small decrease in quality relative to paired T1w segmentation scores (**Fig. S1**).

This adapted SynthSeg pipeline is particularly valuable for studies of neurodevelopment. Previously, commonly used pipelines like SynthSeg and FreeSurfer could not readily be applied across developmental time, necessitating age-specific approaches that led to discontinuities in growth curves (Bethlehem et al., 2022). By extending SynthSeg performance to neonatal MRI scans, this single pipeline can be applied to cohorts from birth to adulthood. Here we reran the new SynthSeg on an aggregated dataset of 26,716 scans, from 0-35 years, showing increases in the number of scans that pass QC until age 10 and preserved performance beyond this. This will prove invaluable for creating lifespan growth charts on large aggregated cohorts, using multiple contrasts without methodological discontinuities. It will further enable dynamic tracking of neurodevelopment in early years, and facilitate the analysis of neurodevelopmental disorders as they present clinically.

Here, we demonstrate how a robust but adult-focussed AI tool for imaging analysis, SynthSeg, fails to generalise to infant scans. This problem is likely pervasive across many emerging segmentation, foundation and clinically-focussed AI models, where younger scans are rarely represented in the training cohorts (Cerri et al., 2026; Gopinath et al., 2025; Henschel et al., 2020; P. Liu et al., 2025). Our proposed mitigation, to rescale infant brain scans, does not require economically and environmentally costly model retraining and is likely to improve performance for other models. However, neonatal brains are not simply rescaled adults. Cortical thickness, sulcal depth and subcortical grey matter, among many other imaging characteristics do not follow a single common growth trajectory, and this complexity likely limits the current approach. In future, deep learning tools trained with data capturing the full neurodevelopmental range will likely improve the imaging of neurodevelopment.

## 5. Conclusions

We have developed and made available an adapted *SynthSeg rescale+crop* pipeline for automated segmentation of neuroanatomical regions across the neurodevelopmental range. We evaluated performance of the pipeline using expertly annotated scans, demonstrating significant improvements in segmentation quality for neonatal MRI scans where *SynthSeg original* consistently failed. Importantly automated QC scores showed corresponding improvements but continue to highlight poor quality segmentations, enabling the application of the new pipeline to large-scale cohorts spanning neurodevelopmental time.

## Data and code availability

We have made the pipeline we developed available at: https://github.com/imaginelab/SynthSeg-resize-and-crop. Code and data to replicate the figures in this manuscript are available at: https://github.com/imaginelab/synthseg-for-developing-brains.

All studies included in the aggregated Neurodevelopmental dataset are open source data, available through different platforms as follows. Multiple datasets are available through different permission groups on the the National Institute of Mental Health (NIMH) Data Archive (NDA): NDA NIMH permissions apply to NIPHD (https://dx.doi.org/10.15154/7m55-b997), IBIS (https://dx.doi.org/10.15154/m84b-k079); UCSD (10.15154/x9wx-cq68); NDA CCF permissions grant access to BCP (https://dx.doi.org/10.15154/nshn-2b72) and HCP-D (https://dx.doi.org/10.15154/21be-hb51); The dHCP data are available through NIMH dHCP permissions (https://doi.org/10.15154/92vw-g837). The ABCD and HBCD data are available on the NBDC Data Hub (https://www.nbdc-datahub.org). The HCP data are available on Connectome DB (https://db.humanconnectome.org/). OpenNeuro hosts the Pixar (https://doi.org/10.18112/openneuro.ds000228.v1.0.0) and Wang (https://doi.org/10.18112/openneuro.ds003604.v1.0.7). The NSPN study is available at: https://nspn.org.uk. The Reproducible Brain Charts Collection ((Shafiei et al., 2025); https://reprobrainchart.github.io) includes: BHRC, devCCNP, PNC.

The BOBS data are openly available on OpenNeuro (https://openneuro.org): https://doi.org/10.17605/OSF.IO/WDR78.

## Author Contributions

**Lena Dorfschmidt:** Conceptualisation, Methodology, Data curation, Software, Visualisation, Writing - Original draft preparation. **Milly Hang Chi Mak**: Data curation, Visualisation, Validation. **Sophie Adler**: Conceptualisation, Supervision, Writing - Original draft preparation, Review & Editing. **Konrad Wagstyl:** Conceptualisation, Supervision, Writing- Original draft preparation, Reviewing and Editing, Funding acquisition.

