## Supplementary figures for "AI segmentation requires accounting for brain size to maintain performance on developmental MRI cohorts"

### Supplementary Materials

| Region | T-statistic | P-value | Bonferroni-corrected p-value |
|---|---|---|---|
| General grey matter | 13.06 | 1.20E-18 | 8.50E-18 |
| General white matter | 11.61 | 1.60E-16 | 1.10E-15 |
| Hippocampus & amygdala | 3.28 | 0.002 | 0.012 |
| Putamen & pallidum | -3.45 | 0.001 | 0.007 |
| Brainstem | 2.91 | 0.005 | 0.036 |
| Cerebellum | 2.58 | 0.013 | 0.089 |
| Thalamus | 0.98 | 0.331 | 1 |

**Table S1.** Statistical comparison of *SynthSeg original* and *rescale + crop* Dice scores based on the 7 brain regions used by SynthSeg for quality control. Using paired t-tests, the largest and most significant improvements were found in the grey and white matter segmentations. Only the putamen and pallidum region had slightly lower Dice scores with rescale+crop

| Region | T-statistic | P-value | Bonferroni-corrected p-value |
|---|---|---|---|
| 3rd Ventricle | 3.45 | 0.001 | 0.032 |
| 4th Ventricle | 3.52 | 8.50E-04 | 0.026 |
| Brain Stem | 2.91 | 0.005 | 0.156 |
| Left Accumbens area | 2.9 | 0.005 | 0.158 |
| Left Amygdala | 3.77 | 3.90E-04 | 0.012 |
| Left Caudate | 4.42 | 4.50E-05 | 0.001 |
| Left Cerebellum Cortex | 2.38 | 0.021 | 0.625 |
| Left Cerebellum White Matter | 1.47 | 0.147 | 1 |
| Left Cerebral Cortex | 8.68 | 6.00E-12 | 1.80E-10 |
| Left Cerebral White Matter | 10.73 | 3.40E-15 | 1.00E-13 |
| Left Hippocampus | 1.19 | 0.24 | 1 |
| Left Inf Lat Vent | 0.55 | 0.585 | 1 |
| Left Lateral Ventricle | 7.34 | 9.60E-10 | 2.90E-08 |
| Left Pallidum | 1.56 | 0.124 | 1 |
| Left Putamen | -1.78 | 0.081 | 1 |
| Left Thalamus | 0.08 | 0.939 | 1 |
| Right Accumbens area | 3.23 | 0.002 | 0.062 |
| Right Amygdala | 0.66 | 0.51 | 1 |
| Right Caudate | 4.41 | 4.70E-05 | 0.001 |
| Right Cerebellum Cortex | 2.58 | 0.013 | 0.377 |
| Right Cerebellum White Matter | 2.71 | 0.009 | 0.264 |

| | | | |
|---|---|---|---|
| Right Cerebral Cortex | 8.25 | 3.00E-11 | 9.10E-10 |
| Right Cerebral White Matter | 12.59 | 5.70E-18 | 1.70E-16 |
| Right Hippocampus | 4.28 | 7.40E-05 | 0.002 |
| Right Inf Lat Vent | 6.49 | 2.40E-08 | 7.20E-07 |
| Right Lateral Ventricle | 5.74 | 4.00E-07 | 1.20E-05 |
| Right Pallidum | 1.38 | 0.174 | 1 |
| Right Putamen | -6.67 | 1.20E-08 | 3.60E-07 |
| Right Thalamus | 1.8 | 0.077 | 1 |
| Right VentralDC | 1.68 | 0.099 | 1 |

**Table S2.** Statistical comparison of *SynthSeg original* and *rescale + crop* Dice scores based for 34 segmented regions. Consistent with Table S1, we found largest improvements in the left and right cerebral cortex and cerebral white matter. We found relatively higher Dice in the pallidum , and that reductions in Dice were only found in the putamen.

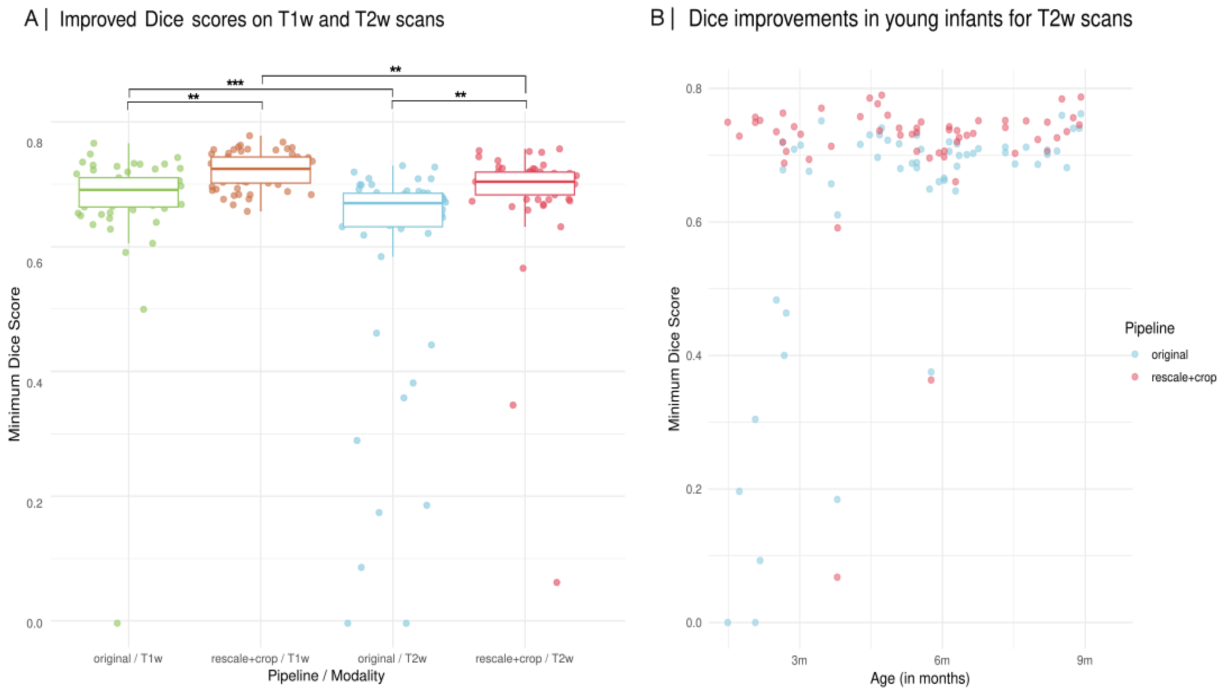


**Figure S1. Testing the new pipeline on T2w scans**

**(A)** We ran both the original and the new pipeline on T2w scans in the BOBs sample. We find that although both pipelines perform better in T1w than T2w scans, the new pipeline still significantly improves Dice scores in T2w scans compared to the original pipeline.

**(B)** With T2w scans, the new pipeline particularly improves Dice scores in younger scans, similar to our results with T1w scans.

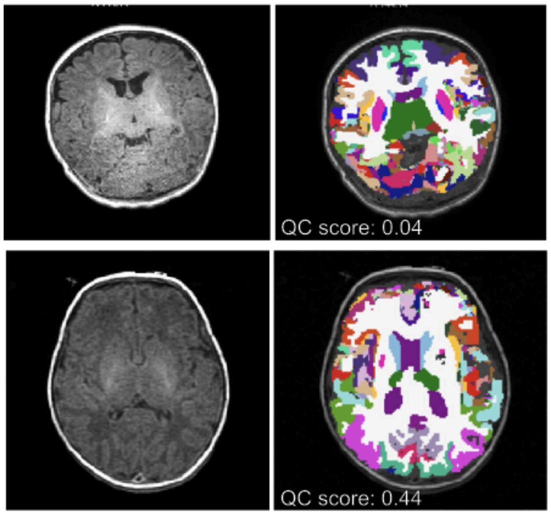


**Figure S2. Example scans with low QC scores from *SynthSeg rescale + crop*.** Visual inspection of example scans with low QC score indicates they are being correctly flagged, supporting the validity of the QC score in the *rescale + crop* pipeline.
